# The Dual Mechanisms of Cognitive Control dataset, a theoretically-guided within-subject task fMRI battery

**DOI:** 10.1101/2021.05.28.446178

**Authors:** Joset A. Etzel, Rachel E. Brough, Michael C. Freund, Alexander Kizhner, Yanli Lin, Matthew F. Singh, Rongxiang Tang, Allison Tay, Anxu Wang, Todd S. Braver

## Abstract

Cognitive control is a critical higher mental function, which is subject to considerable individual variation, and is impaired in a range of mental health disorders. We describe here the initial release of Dual Mechanisms of Cognitive Control (DMCC) project data, the DMCC55B dataset, with 55 healthy unrelated young adult participants. Each participant performed four well-established cognitive control tasks (AX-CPT, Cued Task-Switching, Sternberg Working Memory, and Stroop) while undergoing functional MRI scanning. The dataset includes a range of state and trait self-report questionnaires, as well as behavioural tasks assessing individual differences in cognitive ability. The DMCC project is on-going and features additional components (e.g., related participants, manipulations of cognitive control mode, resting state fMRI, longitudinal testing) that will be publicly released following study completion. This DMCC55B subset is released early with the aim of encouraging wider use and greater benefit to the scientific community. The DMCC55B dataset is suitable for benchmarking and methods exploration, as well as analyses of task performance and individual differences.

## Background & Summary

The Dual Mechanisms of Cognitive Control (DMCC) project was initiated to provide an intensive study of the neural mechanisms associated with cognitive control, a higher mental function utilized to regulate thoughts, emotions, and actions in a flexible and goal-directed fashion^1–4^. The project was designed to test the Dual Mechanisms of Control theoretical framework, which postulates two important modes of control: proactive and reactive^5^. The proactive mode is sustained and anticipatory, with actively maintained goal representations providing an on-going source of bias and top-down input that is used to configure attention and action systems in advance of expected high-demand situations. In contrast, the reactive mode is transient and stimulus-triggered, with conflict or conflict-associated event features activating just-in-time retrieval of goal-relevant information to facilitate task processing. Thus, the Dual Mechanisms of Control framework is a unifying and coherent theoretical account that explains three empirically observed sources of variation: within-individual (task and state-related), between-individual (trait-related), and between-groups (i.e., impaired populations with changes to brain function and integrity), in terms of an underlying dimension involving the temporal dynamics of cognitive control.

The DMCC project is meant to provide a systematic and comprehensive test of the Dual Mechanisms of Control framework. Its goal is to collect both fMRI neuroimaging and behavioral data on a large enough sample for well-powered individual difference analyses. Each participant performs a battery of well-established cognitive control tasks (AX-CPT, Cued task-switching, Sternberg Working Memory, Stroop) while undergoing fMRI scanning, not only under baseline conditions, but also under conditions designed to encourage proactive or reactive control. Each participant takes part in three fMRI scanning sessions (baseline, proactive, reactive), as well as a separate out-of-scanner behavioral session. The behavioral session enables collection of a wide range of individual difference data on psychological health and well-being, personality, and cognitive ability; measures that can be related to the brain imaging data. Longitudinal data, in the form of retesting waves of all sessions, usually spaced more than three months apart, are collected on a subset of participants. Finally, the participant sample includes both a subset of monozygotic (identical) twins, and individuals who previously participated in the Human Connectome Project.

The DMCC project^4^ is currently ongoing, with the data collection effort spanning years, and a target enrollment of N=200. The DMCC55B dataset is the first public release of the DMCC project. The entire DMCC dataset will be released in the future; this subset has been released early to be immediately useful to the wider community, as it already has already proven to be^6,7^. Additional details, particularly of the Proactive and Reactive session task manipulations, are available elsewhere^4^.

The “DMCC55B” name summarizes the data contained in the subset described here: 55 unrelated people, Baseline session only. Although the DMCC55B release is only a subset of the full DMCC, it contains more data on more participants than many fMRI datasets (on a higher “iso-hour contour”^8^), enabling in-depth analysis of individual characteristics. Further, the tasks were presented in a mixed block/event task design^9,10^, which enables a variety of task fMRI modelling options. The functional images were acquired with moderately-high spatial and temporal resolution, using a multiband acquisition sequence. In addition to the task fMRI images, the DMCC55B dataset includes trial-level task performance, structural brain images, physiological monitoring data, and responses on 28 of the individual differences measures collected during the behavioral session. Of the 55 participants included in DMCC55B, 31 were previously participants in the HCP Young Adult study^11^, enabling researchers with HCP data access to combine data from the two studies. The DMCC55B can also be used as a benchmark dataset^12–14^, particularly for methodological investigations focused on higher cognitive brain functions or regions.

## Methods

### Participants

The DMCC55B dataset contains task fMRI and anatomical data from 55 unrelated participants (Mean age=31.7 years, SD=5.9; 34 female). Their self-reported demographic groupings are 39 White/Caucasian; 9 Black/African-American; 4 Asian/Pacific Islander; 2 More Than One Race; 1 Unknown/Prefer Not To Answer. Of the participants, 31 also took part in the HCP Young Adult study^11^. The mapping of DMCC to HCP IDs is available in the HCP ConnectomeDB (db.humanconnectome.org), titled “DMCC (Dual Mechanisms of Cognitive Control) subject key”.

All participants were screened and excluded for MRI safety contraindications. Participants were included if they were between 18 and 45 years of age, without severe mental illness or neurological trauma, and with restricted drug/medication usage (full consent for phone screen^15^). Some participants recruited from the HCP sample would have been excluded under these criteria (e.g., for medical history), but were included if this characteristic was unchanged from the time of their last HCP scan. Participants provided written informed consent in accordance with the Institutional Review Board at Washington University in St. Louis, and received $450 compensation for completion of all sessions, plus a $10 maximum task performance bonus.

Participants were selected from the larger DMCC project for inclusion in DMCC55B by a combination of criteria, with the goal of creating the largest possible (as of December 2020) group of unrelated people with complete and acceptable quality (e.g., minimal overt head motion, consistent task responses) Baseline session data. When both members of a twin pair passed this initial screening, the twin included in DMCC55B was the one subjectively judged to have the highest quality images, specifically the appearance of their quality control images (e.g., clarity of the functional runs’ mean and standard deviation images) and positive control analyses (e.g., distinctiveness of motor region activation to button pressing). In the future, data for all DMCC participants and sessions will be released via the NIMH Data Archive. The 55 participants were chosen from a pool of 97, of whom approximately two thirds were members of twin pairs. The initial screening excluded eleven of the 97 participants for a missing scanning run, six for poor task performance (not responding to five sequential or 40% of the trials in a block), eight for excessive motion, and three for missing Stroop audio recordings (several participants were in more than one of these categories). Twenty twin pairs (40 people) were left in the pool after screening and reviewed individually, with one of each pair selected for final inclusion.

### Behavioral Session

All participants completed an out-of-scanner behavioral session, consisting of task training (illustrated instructions and practice trials), scanner familiarization (if needed), and multiple individual difference instruments. Tasks and questionnaires completed in the laboratory during the behavioral session are indicated by (lab) in the Administration column of Table 1; participants completed the others either at home or during a later session in the laboratory, as they preferred.

**Table 1.**
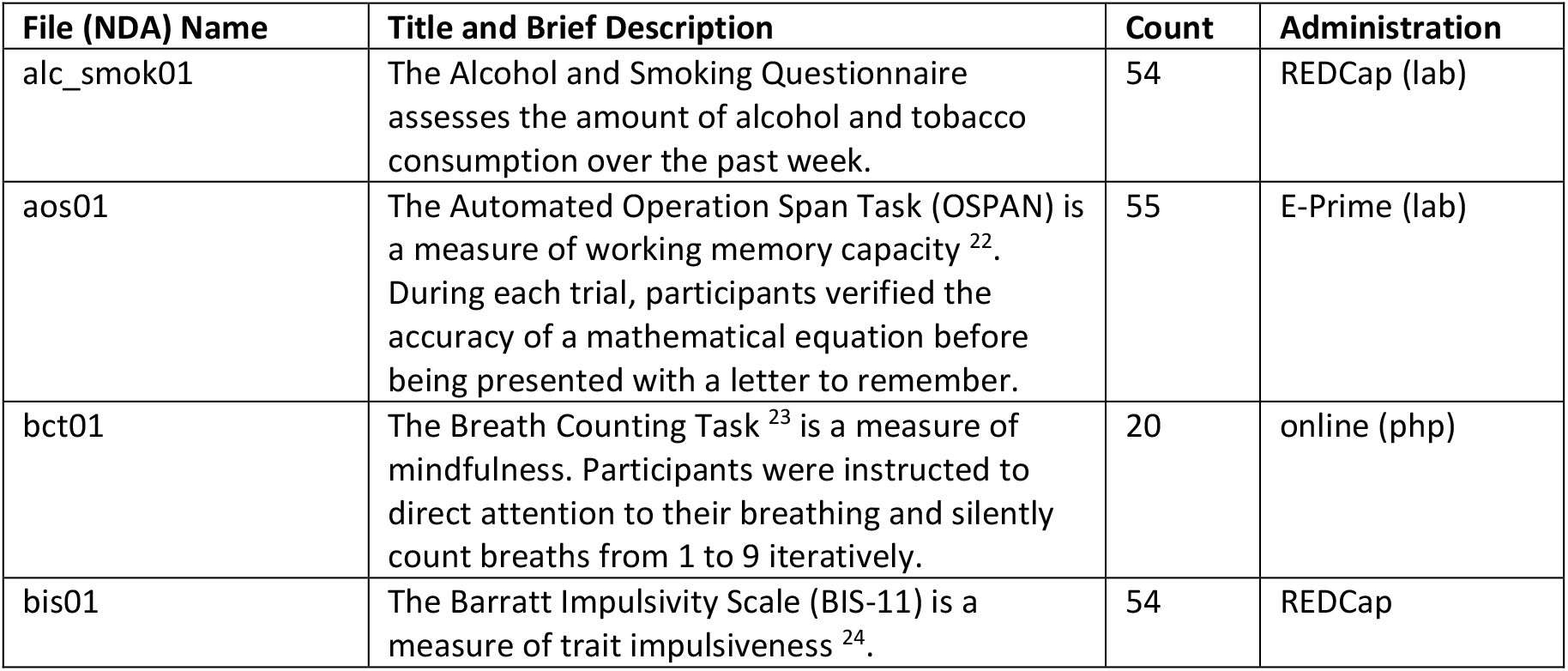
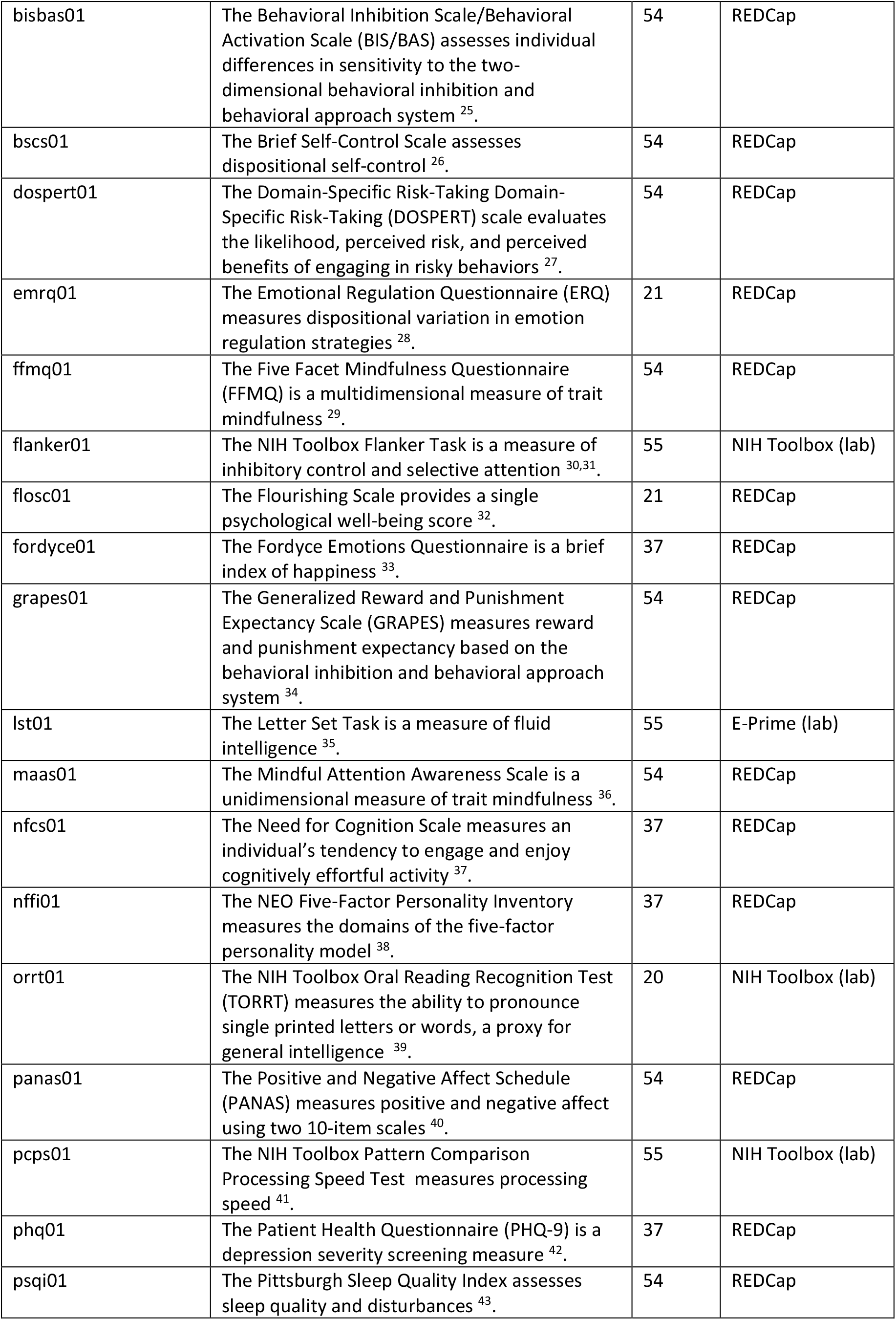
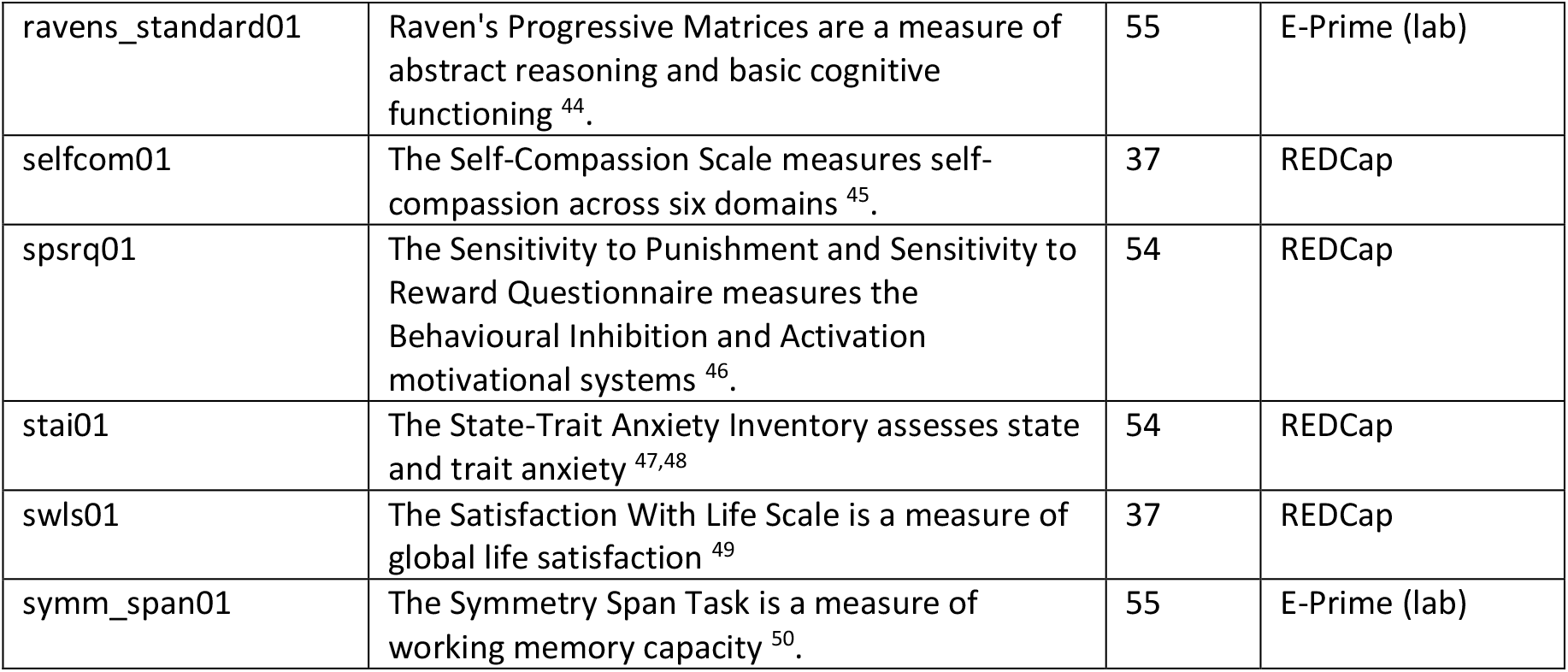
DMCC55B questionnaire data. Scores for each assessment are in a separate derivatives^16^ file, named according to its NDA data dictionary definition, as listed in “File (NDA) Name”. “Count” lists the number of participants with data for each questionnaire, which varies since one participant did not complete any REDCap questionnaires and some questionnaires were added over time. behavioralSession_individualDifferences^18^ has group and individual performance summary statistics for these measures. Administration was by REDCap^19,20^, the NIH Toolbox^21^, E-Prime (2.0, RRID:SCR_009567), or custom online php scripts, as indicated.

The assessments released in DMCC55B (Table 1) span personality and psychological health and well-being, cognitive tests of crystallized and fluid intelligence, processing speed, working memory capacity, and attentional control. Scores and responses for each participant are available under derivatives in the OpenNeuro dataset^16^, in files formatted and named according to the NIMH Data Archive (NDA^17^) specifications, with the exception of certain restricted information (GUIDs, demographics), which was omitted for privacy. All DMCC questionnaire data will be made public through the NDA in the future. In addition to individual response files, group averages and derived scores are located in the file behavioralSession_individualDifferences^18^.

### Imaging Session

All scans were acquired using a 3T Siemens Prisma with a 32-channel head coil in the East Building MR Facility of the Washington University Medical Center. Functional (BOLD, blood oxygenation level dependent) scans were acquired with CMRR multiband sequences (University of Minnesota Center for Magnetic Resonance Research)^51–53^, multiband (simultaneous multislice; SMS) factor 4, without in-plane acceleration (iPat = none), resulting in 2.4 mm isotropic voxels, 1.2 s TR (TE 33 msec, flip angle 63°; protocol sheets at^15^). All functional runs were collected with both Anterior to Posterior and Posterior to Anterior encoding directions (“AP” and “PA”, respectively); the first run of each task was always AP, the second immediately followed and was PA. One pair (AP, PA) of spin echo field map images was collected before the first functional scan, and again upon re-entry if the participant left the scanner during the session. A single-band reference (“SBRef”) image with matching acquisition parameters was collected immediately before every functional run. T1 and T2-weighted high-resolution MPRAGE structural scans were acquired at the beginning of the scanning session (T1: 0.8 mm isotropic voxels; 2.4 s TR, 0.00222 s TE, 1 s TI, 8° flip angle; T2: 0.8 mm isotropic voxels; 3.2 s TR, 0.563 s TE, 120° flip angle). All files were automatically sent from the scanner to an XNAT (RRID:SCR_003048) database managed by the Connectome Coordination Facility (CCF, https://www.humanconnectome.org/)^54^ for permanent archiving and initial data verification.

Finger photoplethysmograph and respiration belt recordings during scanning were made with Siemens equipment. The eye tracking system camera was positioned to allow research technicians to monitor participant wakefulness from within the control room, but no recordings were made nor eye tracking performed. Additionally, since February 2018, the FIRMM (https://firmm.readthedocs.io/en/3.2/)^55^ system was used for participant motion monitoring; run duration was not varied based on FIRMM output, but participants were informed and/or repositioned if FIRMM indicated excessive movement (FIRMM present for 23 of the 55 participants). Stroop responses were verbal and recorded via microphone. At the beginning of data collection a Micro-Optics^56^ microphone was used for this purpose, but in November 2017 a switch was made to the FOMRI-III^57^ microphone, due to mechanical failure of the original system.

The Baseline imaging session typically began with the anatomical scans, followed by the first resting state run (5 minutes), then the first two tasks (four runs total; two of each task), then the second resting state run (also 5 minutes; resting state scans are not included in this data release), and finally the remaining two pairs of task runs. Due to time constraints, resting state runs were occasionally moved to the end of the session or omitted. If the Baseline structural scans were not rated of sufficient quality (structural rating scale and criteria following HCP protocols) they were repeated at the beginning of a later scanning session and the highest-rated scans were used (12 DMCC55B people include anatomy from a later session).

## Tasks

The full DMCC protocol contains four cognitive control tasks, each of which are performed in three separate scanning sessions: Baseline, Proactive, and Reactive. The task details (instructions, stimuli, etc.) varied across sessions to encourage the use of proactive or reactive cognitive control^4^. The DMCC55B dataset contains Baseline session task runs only, so the Baseline version of each task is described here.

The tasks were presented via Eprime (2.0, RRID:SCR_009567, Psychology Software Tools, Pittsburgh, PA); scripts are available for download^58^. A csv version of the Eprime output for each task run is included in the OpenNeuro derivatives^16^. Code for extracting and calculating the behavioral and timing measures used for analysis is in the dualmechansisms github site and the TaskPerformance^18^ results summary files for each task. In all tasks except Stroop, task responses were button presses with the index or middle finger of the right (always) hand; Stroop responses were verbal.

Each task run followed a mixed, block/event-related format^9,10^: trials were not evenly spaced throughout the run but rather grouped into three long blocks. Each task block lasted approximately 3 minutes and was preceded and followed by a 30 second fixation block, for runs of approximately 12 minutes (24 minutes total per task). The precise trial and run durations varied slightly between tasks; see below and DMCC_taskTrialSummary.docx^15^. Trial duration and inter-trial intervals (ITIs) were timed to synchronize with the scanner pulses to facilitate estimating event-related hemodynamic responses.

The four tasks are described next, in alphabetical order (participants completed the tasks in random order).

### AX-CPT (Axcpt)

In the DMCC version of the AX-CPT task^59–62^ participants are asked to respond to two-event trials (Figure 1). Each trial begins with a Cue (always a letter), followed by a delay of 4.3 seconds, and ends with a Probe, which can be a letter or a number. The DMCC version comprises six types of trials; the type, number (total, not per run), and an example of each are listed in Figure 1. The same stimuli (letters and numbers) are used for all participants and listed in the Eprime output; trial order is random.

**Figure 1.**
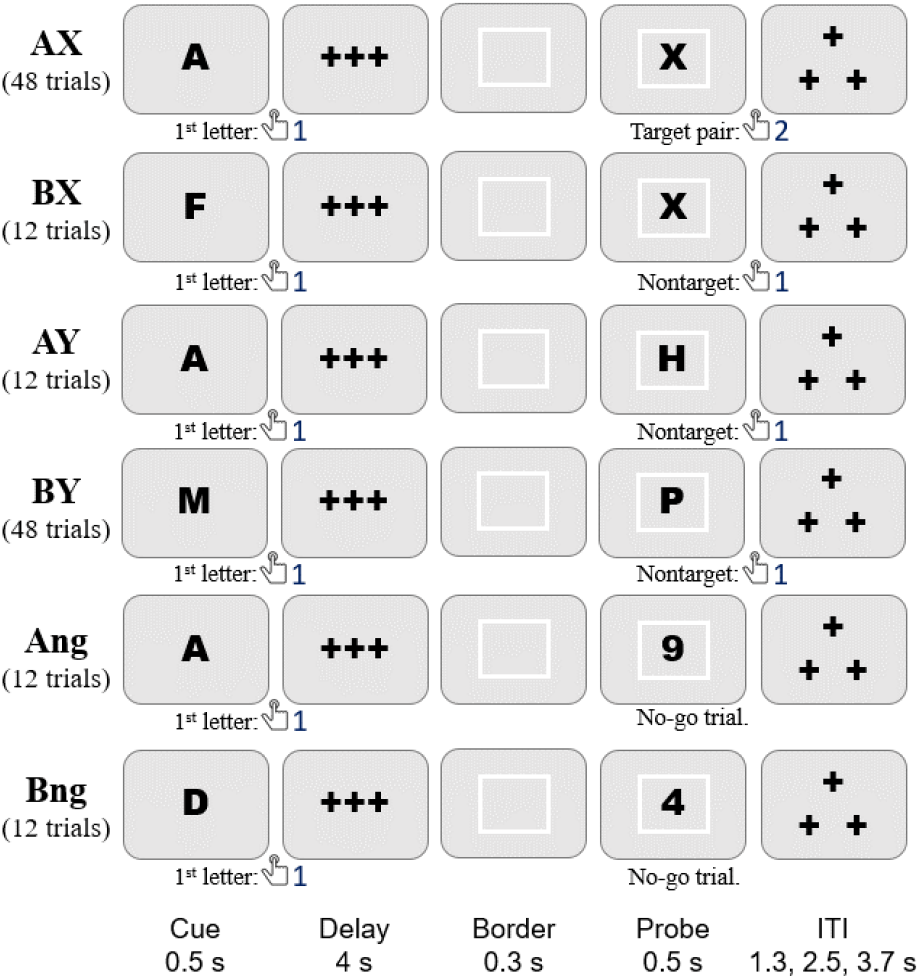
AX-CPT task timing, trial types, and correct responses. The total number of trials of each type completed by each participant is listed. Task instructions and training materials in AXCPT_Cards.pptx^15^; scoring code and DMCC55B summary in taskPerformance_Axcpt^18^.

**Figure 2.**
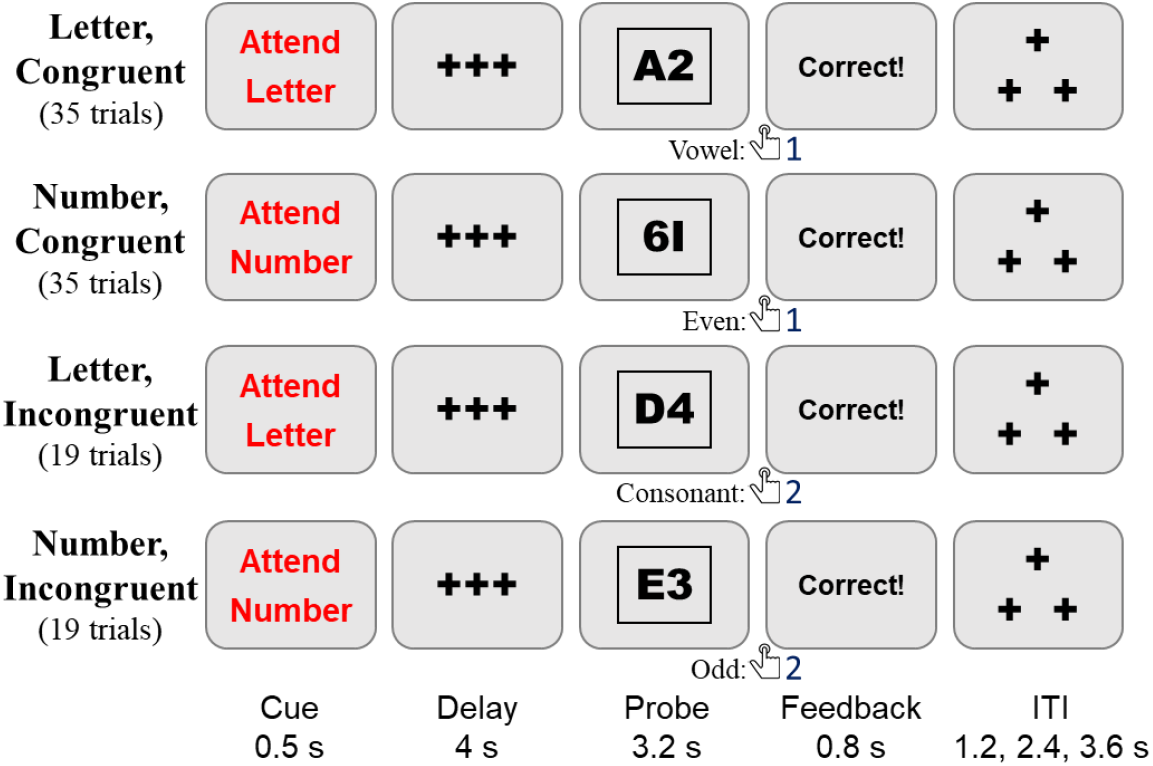
Cued task-switching task timing, trial types, and correct responses. The total number of trials of each type completed by each participant is listed. Task instructions and training materials in CuedTS_Cards.pptx^15^; scoring code and DMCC55B summary in taskPerformance_Cuedts^18^.

The AX-CPT task requires two responses per trial: the first (always button 1) after the Cue, and the second (button 2 if a target pair, button 1 for nontarget, and none if a no-go trial) at the Probe. The AX trial type (Cue A Probe X) is the target; these are only trials for which the correct Probe response is button 2. No-go trials (Ang, Bng) are indicated by a number Probe (rather than letter); withholding a button press is the correct response. All other trial types (AY, BX, BY) are nontarget, with button 1 as the correct Probe response. The response_time, response_button, and response_accuracy columns of the _events.tsv BIDS files^16^ correspond to the Probe response; Cue responses are only in the Eprime output.

Cue responses are expected to have very low error rates (and fast reaction times), as the correct response is always button 1. For Probe responses, performance is expected to vary with trial type: error rate higher and RT slower on AY and BX trials than AX and BY trials. In DMCC analyses, BX trials were considered to have high cognitive control demands (because of the response conflict occurring with the Probe, which required use of the Cue for context), while BY trials have low control demands (because the Probe is always associated with a button 1 response).

### Cued Task-Switching (Cuedts)

The DMCC uses as letter-digit version of the Cued task-switching task^63–65^: the Probe always consists of a single number and letter. The Cue instructs the participant to either “Attend Letter” (respond with button 1 if the letter is a vowel; button 2 if a consonant) or “Attend Number” (respond with button 1 if the number is even; button 2 if odd). Since each Probe has both a letter and number, and the response to both Cues is one of two button presses, the stimuli can be further described as Congruent or Incongruent. On Congruent trials, the same response should be made to the Probe, irrespective of the Cue; conversely, on Incongruent trials, the correct response to the Probe depends on the Cue (i.e., what task was instructed). For example, the Probe 1B is Congruent: an odd number (button 2) and a consonant letter (button 2), so button 2 is always the correct response. Likewise, Probe 2B is Incongruent: if the Cue was Attend Number, then the correct response is button 1 (even); if the Cue was Attend Letter, then the correct response is button 2 (consonant).

The trials can further be described as repeat (when the current trial Cue is the same as that of the previous trial) or switch (when the current trial Cue is different than that of the previous trial). The trial_type (Congruent, Incongruent), trial_cue (AttendLetter, AttendNumber), and trial_switch (switch, repeat) are listed in the _events.tsv files; the stimulus for each trial is in the Eprime output. The same trials (Cue-Probe pairs) are presented to each participant, but their order within blocks is random, so the number of switch and repeat trials varies somewhat between participants (approximately half of the trials are switch and half repeat, but the first trial of every block is neither, causing slight differences in counts across participants).

In DMCC analyses, Incongruent trials were considered to have high cognitive control demands (because of the importance of the Cue and the response conflict occurring with the Probe), while Congruent trials have low control demands (because the Probe response does not depend on the Cue). Incongruent trials were expected to have generally higher error rates and slower reaction times relative to Congruent trials; a similar pattern was expected on switch relative to repeat trials.

### Sternberg Working Memory (Stern)

The DMCC version of the Sternberg task^66–68^ uses lists with 5 to 8 words (Figure 3; trial_length column of _events.tsv^16^), split between two List screens, followed (after a 4 second Delay) by a Probe word. Participants are asked to press button 2 if the Probe was in either List of the current trial, and button 1 if it was not. The Probe word was a member of the List in half of the trials (45 total), with this trial type termed NP (“novel positive”). In most of the remaining trials (36 total) the Probe is new, not in any previous List (type NN, “novel negative”). However, in 9 trials the Probe was in the *immediately preceding* trial’s List (type RN, “recent negative”). All participants were given the same words, divided into the same Lists, and in the same order (SternbergWordKey.txt on the dualmechansisms github site^69^). The trial type frequencies varied according to List length, with five-word Lists having the highest frequency, as these were the focus of comparisons with the Proactive and Reactive conditions.

**Figure 3.**
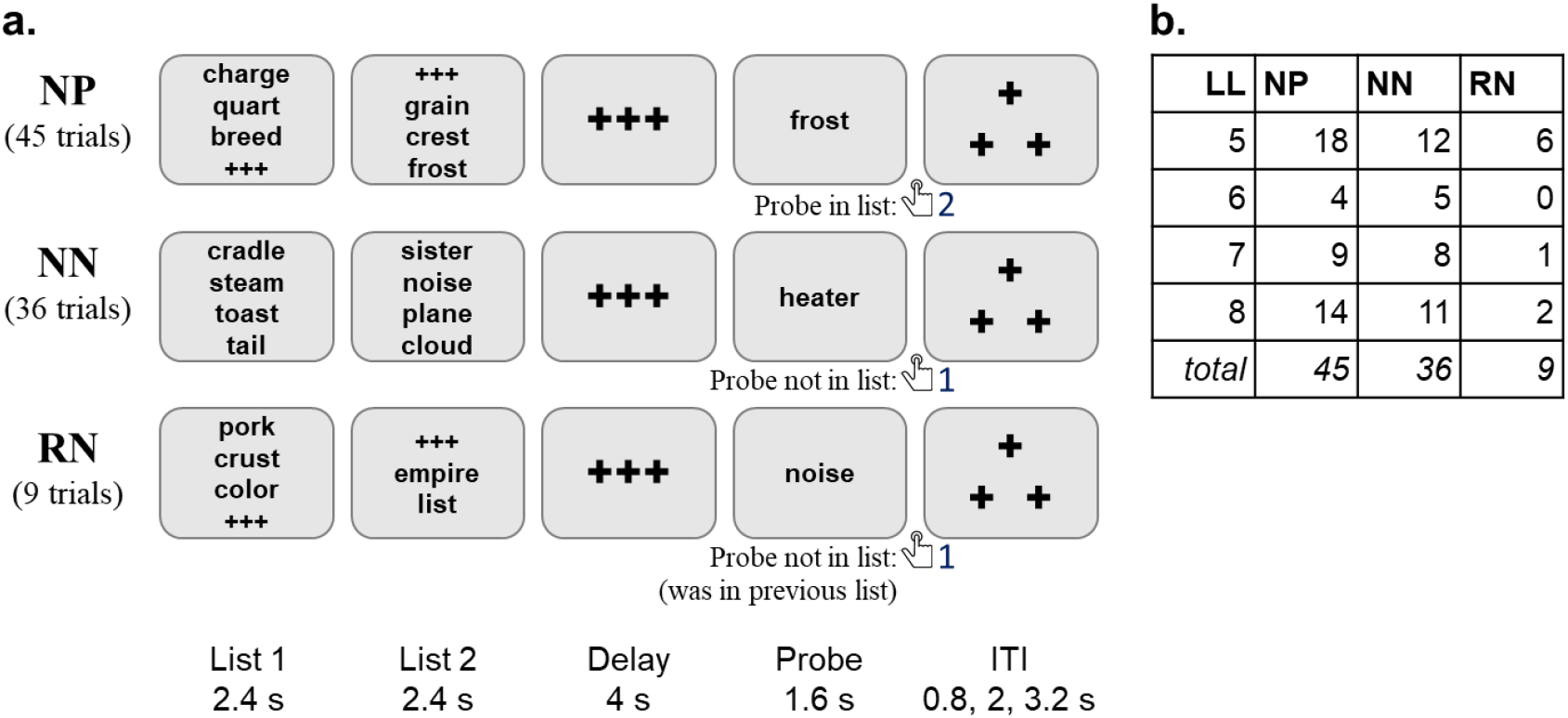
Panel a: Sternberg task timing, trial types, and correct responses. The total number of trials of each type completed by each participant is listed. Note that there is an implied top-to-bottom sequential order in these example trials: the third example is of type RN because its Probe word (“noise”) was in the previous trial’s word List 2. The ListLength (LL) attribute is the number of words in List 1 and List 2 combined: the first (NP) example has length 6, second (NN) length 8, and third (RN) length 5. Panel b: Total number of examples of each type and length. Task instructions and training materials in Sternberg_Cards.pptx^15^; scoring code and DMCC55B summary in taskPerformance_Sternberg^18^.

In DMCC analyses, RN trials were considered to have high cognitive control demands (because the recent familiarity of the Probe would cause interference with processing based on working memory contents), while NN trials were considered to have low control demands (since Probes had no familiarity). The RN trials were expected to have the highest error rates and slowest reaction times, and NN trials the lowest error rate and fastest reaction time. Reaction time and error rate were also expected to increase with List length.

### Color-Word Stroop (Stroop)

The DMCC version of the classic Stroop task^70^ is a color-word variant, with verbal responses (the participant is asked to say the ink color rather than the printed word). The trial types are described in terms of congruency: Congruent if the printed word indicates a color that is the same as the font color (“Con”, two thirds of the trials); Incongruent if the printed word indicates a color that is different from the font color (“InCon”, one third of the trials); Figure 4 (trial_type column of _events.tsv^16^).

**Figure 4.**
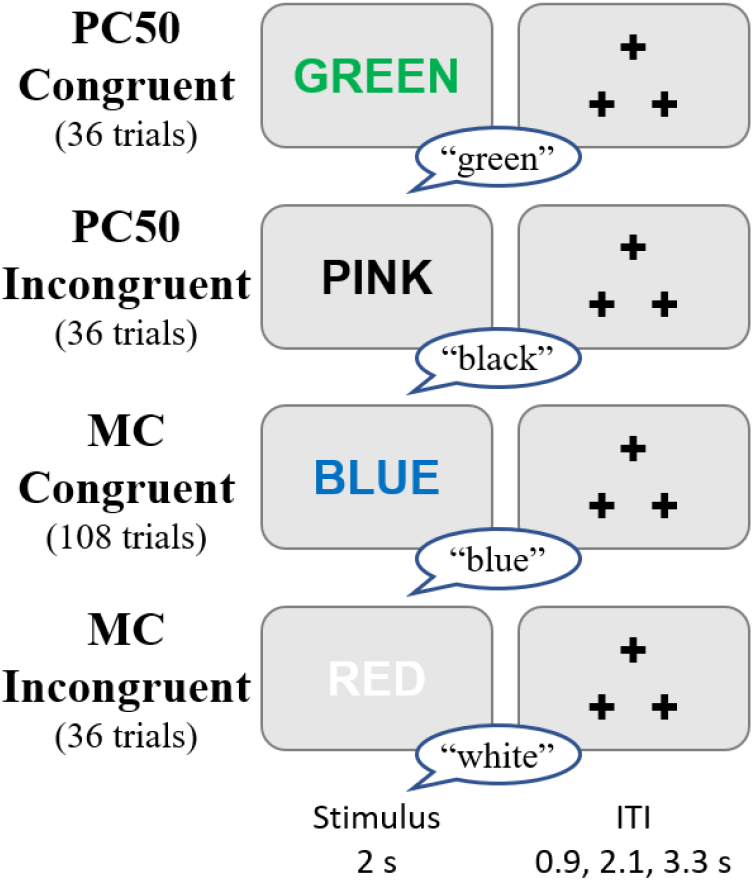
Stroop task timing, trial types, and correct responses. The total number of trials of each type completed by each participant is listed. Task instructions and training materials in Stroop_Cards.pptx^15^; scoring code and DMCC55B summary in taskPerformance_Stroop^18^.

A total of eight colors were used in the DMCC Stroop task, divided into two sets of four colors each (to enable different types of proportion congruency manipulations^71,72^). The first set of colors (red, blue, white, and purple) is termed “MC” (mostly congruent) and includes 108 Congruent and 36 Incongruent trials. The second set (pink, green, black, and yellow) is termed “PC50” (50% percent congruent), and includes 36 Congruent and 36 Incongruent trials (i.e., an equal split). The PC50 trial type was used to facilitate comparisons with the Proactive and Reactive conditions, as they occurred with identical frequency in all three sessions. The color set (PC50 or MC) of each trial is listed in the trial_lwpc column of the _events.tsv file; the colors and words are in the full Eprime output.

In DMCC analyses, Incongruent trials were considered to have high cognitive control demands (because of the response conflict between color and word dimensions) while Congruent trials had low control demands (because both dimensions indicated the same response). Incongruent trials were expected to have slower reaction times and higher error rates than Congruent trials.

## Preprocessing

Preprocessing of fMRI data was performed with fMRIPrep 1.3.2 (RRID:SCR_016216)^73,74^, which is based on Nipype 1.1.9 (RRID:SCR_002502)^75,76^. For implementation, fMRIPrep was run within a Singularity^77^ container on a Linux server managed by the Washington University in St. Louis NeuroImaging Laboratories Computer Support Group. Custom bash and python scripts were created to implement file transfer before and after the fMRIPrep container as well as other parts of the pipeline. The following text describing the anatomical and functional data preprocessing was automatically generated by fMRIprep^73^, and minimally edited for style.

### Anatomical data preprocessing

The T1-weighted (T1w) image was corrected for intensity non-uniformity (INU) with N4BiasFieldCorrection^78^, distributed with ANTs 2.2.0^79^(RRID:SCR_004757), and used as T1w-reference throughout the workflow. The T1w-reference was skull-stripped with a Nipype implementation of the antsBrainExtraction.sh workflow (from ANTs), using OASIS30ANTs as target template. Brain surfaces were reconstructed using recon-all (FreeSurfer 6.0.1, RRID:SCR_001847)^80^, and the brain mask estimated previously was refined with a custom variation of the method to reconcile ANTs and FreeSurfer-derived segmentations of the cortical gray-matter of Mindboggle (RRID:SCR_002438)^81^. Spatial normalization to the ICBM 152 Nonlinear Asymmetrical template version 2009c (RRID:SCR_008796)^82^ was performed through nonlinear registration with antsRegistration (ANTs 2.2.0), using brain-extracted versions of both T1w volume and template. Brain tissue segmentation of cerebrospinal fluid (CSF), white-matter (WM) and gray-matter (GM) was performed on the brain-extracted T1w using fast (FSL 5.0.9, RRID:SCR_002823)^83^.

### Functional data preprocessing

For each of the BOLD runs found per subject), the following preprocessing was performed. First, a reference volume and its skull-stripped version were generated using custom fMRIPrep methodology. A deformation field to correct for susceptibility distortions was estimated based on two echo-planar imaging (EPI) references with opposing phase-encoding directions, using 3dQwarp^84^ (AFNI 20160207). Based on the estimated susceptibility distortion, an unwarped BOLD reference was calculated for a more accurate co-registration with the anatomical reference. The BOLD reference was then co-registered to the T1w reference using bbregister (FreeSurfer) which implements boundary-based registration^85^. Co-registration was configured with nine degrees of freedom to account for distortions remaining in the BOLD reference. Head-motion parameters with respect to the BOLD reference (transformation matrices, and six corresponding rotation and translation parameters) were estimated before spatiotemporal filtering using mcflirt (FSL 5.0.9)^86^. BOLD runs were slice-time corrected using 3dTshift from AFNI 20160207 (RRID:SCR_005927)^84^. The BOLD time-series, were resampled to the fsaverage5 surface. The BOLD time-series (including slice-timing correction) were resampled onto their original, native space by applying a single, composite transform to correct for head-motion and susceptibility distortions. These resampled BOLD time-series will be referred to as preprocessed BOLD in original space, or just preprocessed BOLD. The BOLD time-series were resampled to MNI152NLin2009cAsym standard space, generating a preprocessed BOLD run in MNI152NLin2009cAsym space. A reference volume and its skull-stripped version were generated using custom fMRIPrep methodology.

Several confounding time-series were calculated based on the preprocessed BOLD: framewise displacement (FD), DVARS, and three region-wise global signals. FD and DVARS are calculated for each functional run, both using their implementations in Nipype (following the definitions^87^). Three global signals are extracted: the CSF, the WM, and the whole-brain masks. Additionally, a set of physiological regressors were extracted to allow for component-based noise correction (CompCor)^88^. Principal components are estimated after high-pass filtering the preprocessed BOLD time-series (using a discrete cosine filter with 128s cut-off) for the two CompCor variants: temporal (tCompCor) and anatomical (aCompCor). Six tCompCor components are then calculated from the top 5% variable voxels within a mask covering the subcortical regions. This subcortical mask is obtained by heavily eroding the brain mask, which ensures it does not include cortical GM regions. For aCompCor, six components are calculated within the intersection of the aforementioned mask and the union of CSF and WM masks calculated in T1w space, after their projection to the native space of each functional run (using the inverse BOLD-to-T1w transformation). The head-motion estimates calculated in the correction step were also placed within the corresponding confounds file. All resamplings can be performed with a single interpolation step by composing all the pertinent transformations (i.e. head-motion transform matrices, susceptibility distortion correction when available, and co-registrations to anatomical and template spaces). Gridded (volumetric) resamplings were performed using antsApplyTransforms (ANTs), configured with Lanczos interpolation to minimize the smoothing effects of other kernels^89^. Non-gridded (surface) resamplings were performed using mri_vol2surf (FreeSurfer).

Many internal operations of fMRIPrep use Nilearn 0.5.0 (RRID:SCR_001362)^90^, mostly within the functional processing workflow. For more details of the pipeline, see the section corresponding to workflows in fMRIPrep’s documentation.

### Physiological Recordings

Finger photoplethysmograph and respiration belt recordings were extracted using https://github.com/CMRR-C2P/MB/blob/master/readCMRRPhysio.m and then converted to plain text; no filtering or other processing was performed. All collected recordings (41 participants) are included with DMCC55B, regardless of clarity.

### Stroop Responses

Audio recording (to .wav files) of the verbal response for each Stroop trial was triggered by the Eprime task script. Two research assistants listened to each audio clip and recorded the word they heard; the consensus response was listed and coded for accuracy (response_accuracy column of _events.tsv^16^). Unclear responses were coded as “unintelligible” or “no-response”; we suggest treating unintelligible trials as missings and no-response as incorrect (0). The reaction time in each Stroop audio recording was calculated via in-house MATLAB (RRID:SCR_001622) scripts custom-designed for extracting voice onsets from noisy audio files (audio section of the dualmechanisms github site), and are in the response_time column of _events.tsv. The MATLAB scripts (one version for each microphone) were validated in a subset of recordings by comparing its reaction times to those of two observers listening to the same recordings with Audacity recording and editing software (RRID:SCR_007198) and marking the reaction time.

## Data Records

The DMCC55B data records are organized following the BIDS (Brain Imaging Data Structure) standard (version 1.4.0)^91^, and are available as OpenNeuro Dataset ds003465^16^. Supporting materials such as checklists used during scanning, recruitment scripts, and task instructions are available at The Dual Mechanisms of Cognitive Control Open Science Foundation (OSF) website^15^. DMCC55B-specific material such as performance and quality control summaries are in the Publications, DMCC55B Dataset Description subsection^18^ of the main DMCC OSF site^15^. After 2023, qualified researchers will be able to obtain data via the NIMH Data Archive (nda.nih.gov, collection 2970) as well.

The questionnaire data records are included in the DMCC55B derivatives^16^, named according to the NDA data dictionary entry which defines its format. For example, suppose a researcher is interested in the Positive and Negative Affect Schedule (PANAS) questionnaire scores. In Table 1 its “File (NDA) Name” is panas01, so the participants’ scores are in panas01_DMCC55B.csv^16^, which is defined by https://nda.nih.gov/data_structure.html?short_name=panas01.

## Technical Validation

### Standard Procedures and Reports

A key component of the strategy to ensure accurate and high-quality data collection is the use of Standard Operating Procedures (SOPs) documents and checklists, available in the Sessions and Storage section of the DMCC OSF site^15^. The SOPs describe how to carry out all parts of the DMCC protocol: recruitment, data collection (behavioral and scanning sessions), data storage, preprocessing, quality assessment, and analysis. SOPs are used for training, but more importantly, for daily reference, and updated as needed (e.g., after an equipment change or with new troubleshooting tips). Accompanying paper checklists (digitized for storage) are used during interactions with the participants to reduce the likelihood that a data collection or instructional step is skipped, and to collect notes during the sessions.

Although automated preprocessing and analysis pipelines were employed, manual inspection and QC/QA data review is still essential^92–94^. Accordingly, the pipelines were designed to stop at key points for quality checks, as well as when encountering unexpected data. The DMCC generates standard reports on each participant’s data (primarily as knitr^95^ documents), including behavioral performance, movement, image clarity, physiological recordings, and GLM contrasts. Successful compilation of these reports indicates that the source files are in the expected locations, names, and formats. Importantly, the reports collect key metrics into concise files, enabling relatively quick review after each scanning session, and adjustments if needed before future sessions. Examples of these reports are in the Analyses section of the DMCC OSF site^15^.

### Task Performance

The task performance suggests that participants understood and followed the task instructions. Error rates were generally low and reaction times fast, while keeping below ceiling. As a validation check on the cognitive control demand manipulation, reaction times were confirmed to be slower and error rates higher in the high relative to low control condition (the high-low distributions in Figure 5 show most participants greater than zero in all tasks). Other expected performance patterns were also found, such as longer reaction time and more errors on Switch than Repeated trials in Cued task-switching, and with increasing List length in Sternberg, shown in the taskPerformance files^18^ for each task.

**Figure 5.**
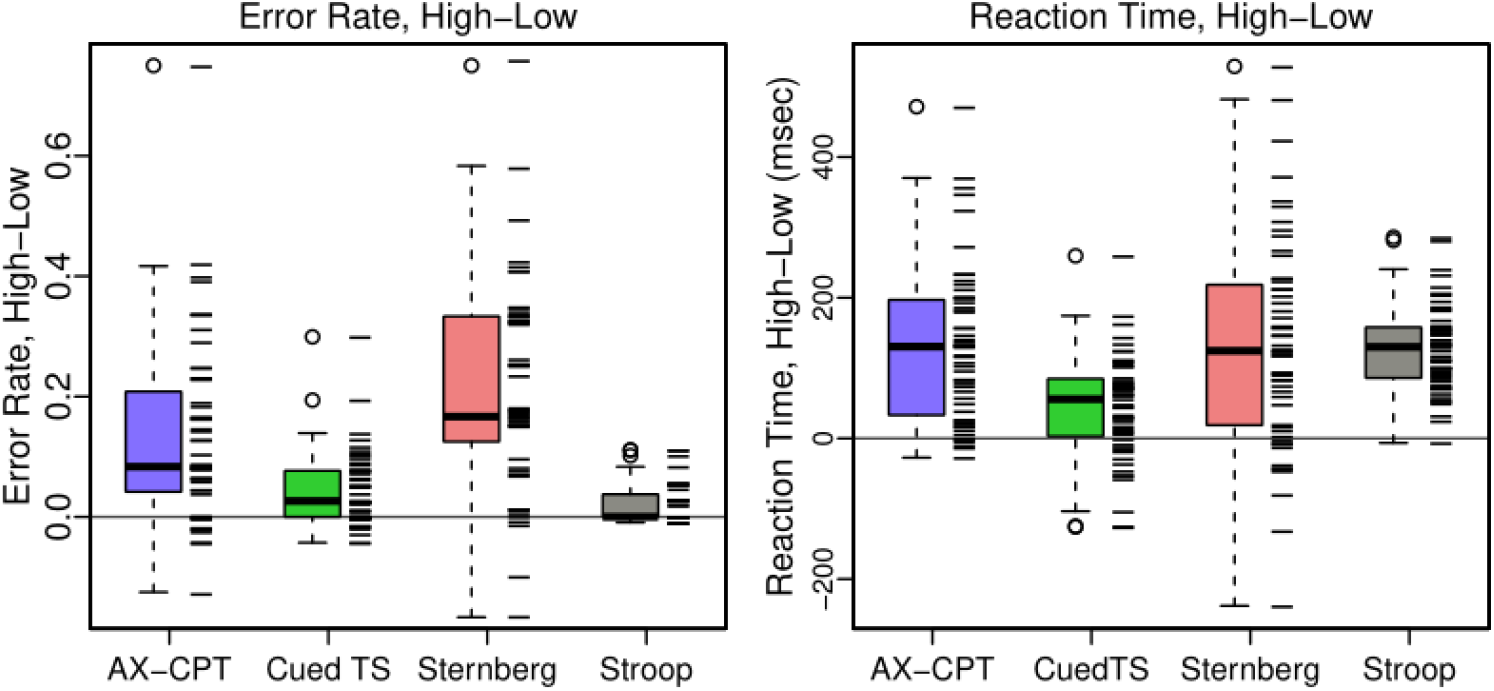
Distribution of high - low cognitive control differences on each task. As expected, both the error rate and reaction time were greater for high than low conditions, and so most differences are positive. Tick marks show the values for each participant, with a small amount of jitter added to the error rate to reduce overplotting. Means for each person and all trial types are in the taskPerformance files^18^.

### Motion (Apparent and Overt)

It is always critical to both assess and minimize motion during MRI scanning. With multiband fMRI scanning (as used here) evaluating head motion can be somewhat more difficult, as the realignment parameters contain a combination of overt and apparent motion^96–98^. Following convention we summarize motion here and set the censoring threshold by Framewise Displacement (FD > 0.9 mm)^99,100^, despite considering the six separate realignment parameters more useful for quality control, as the different types of “motion” are readily distinguishable^101^. Timeseries plots for the six realignment parameters and FD for every participant and run suggest that DMCC55B participants had relatively low motion, both overt and apparent (QC_motion^18^).

The distribution of FD for each participant and task is summarized in Figure 6. These plots suggest that FD varies more with participant than task; some participants have higher FD than others, but the distributions for each task within a participant tend to be similar. The median framewise displacement across subjects is 0.16 mm, seven people have no censoring in any run, and only two participants have more than 5% of the frames above the censoring threshold (in a single task each).

**Figure 6.**
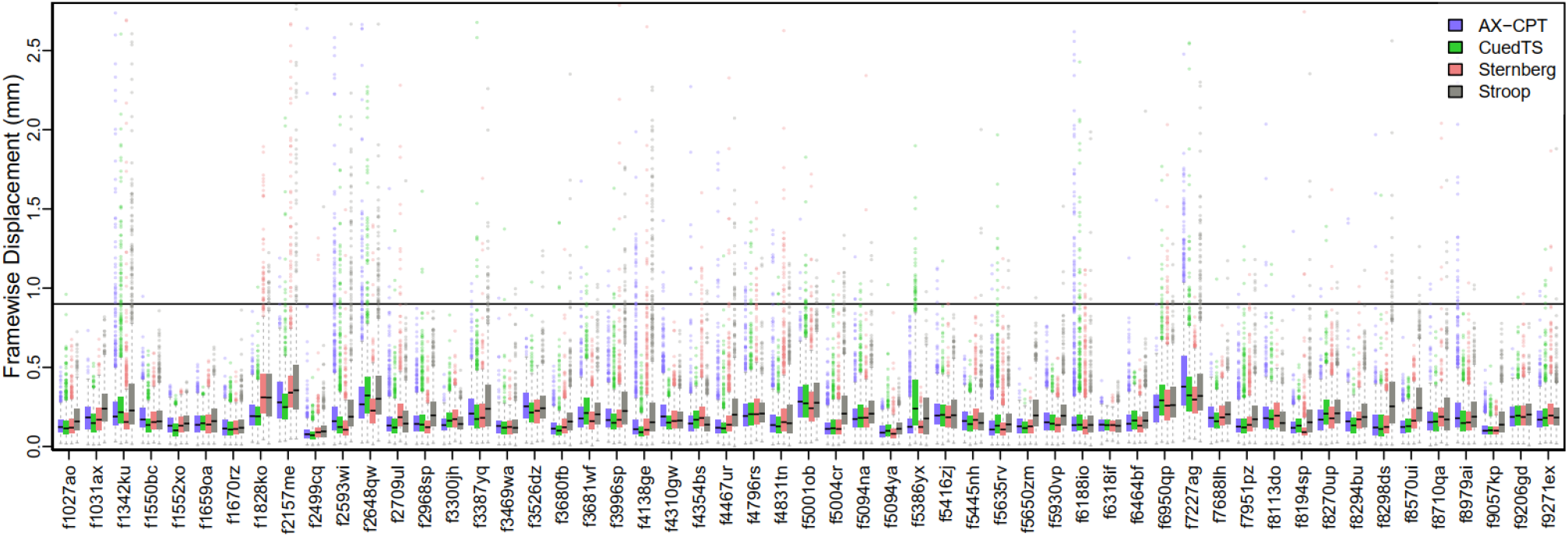
Framewise Displacement for each subject and task (frames from both runs combined for each task). The line at FD 0.9 is the value at which we censor frames for the DMCC GLMs^99,102^. Only two participants had a task with more than 5% of the frames reaching the 0.9 mm FD censoring threshold: f7227ag on AX-CPT and f1342ku on Stroop. The percent of frames censored and median FD for each person is in QC_motion^18^.

### tSNR (temporal Signal to Noise Ratio)

Temporal signal to noise ratio (tSNR) images are calculated by dividing the mean by the standard deviation of all frames within a run in each voxel (or vertex), after preprocessing. While not isolating task signal, tSNR (as well as the temporal mean and standard deviation) images are valuable for identifying acquisition artifacts, areas of signal dropout, and relative differences across participants or runs^103–107^. Figure 7 shows the median volumetric tSNR image. Images of the cortical surface and individual participants are in the QC SD and tSNR files on the DMCC55B OSF site^18^.

**Figure 7.**
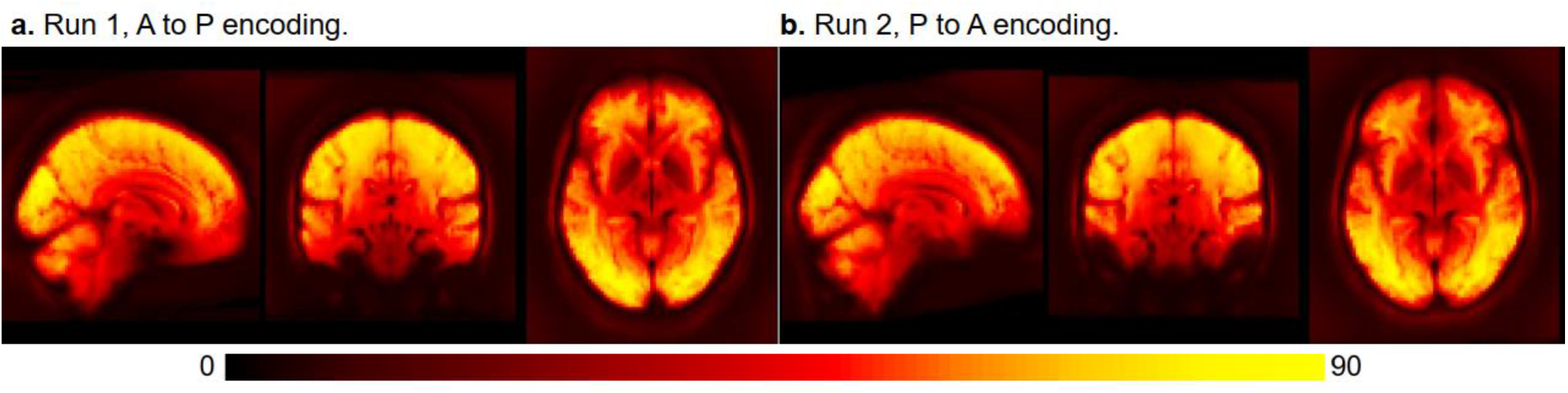
Median tSNR in each voxel. This image is not an individual participant, but rather the median calculated over all people and tasks, each run separately. Details and a surface version are in QC_tSNR_summary^18^.

Anatomical structure is clearly visible in the median DMCC55B tSNR image for each encoding direction (Figure 7), even in subcortical structures, suggesting generally high image quality. As expected, there is some medial frontal and temporal signal dropout near the air-filled sinuses and ear canals, with the most affected area varying with encoding direction (AP or PA). The distribution of tSNR over all vertices for each task and participant is shown in Figure 8. As with FD, most tSNR variance occurs between people (e.g., higher tSNR for f1552xo than f1342ku) rather than for tasks within each person, suggesting that the differing task demands did not systematically affect image quality. The median tSNR of surface vertices across all participants and tasks is 103.68, which compares favorably to other multiband task fMRI datasets^108–110^.

**Figure 8.**
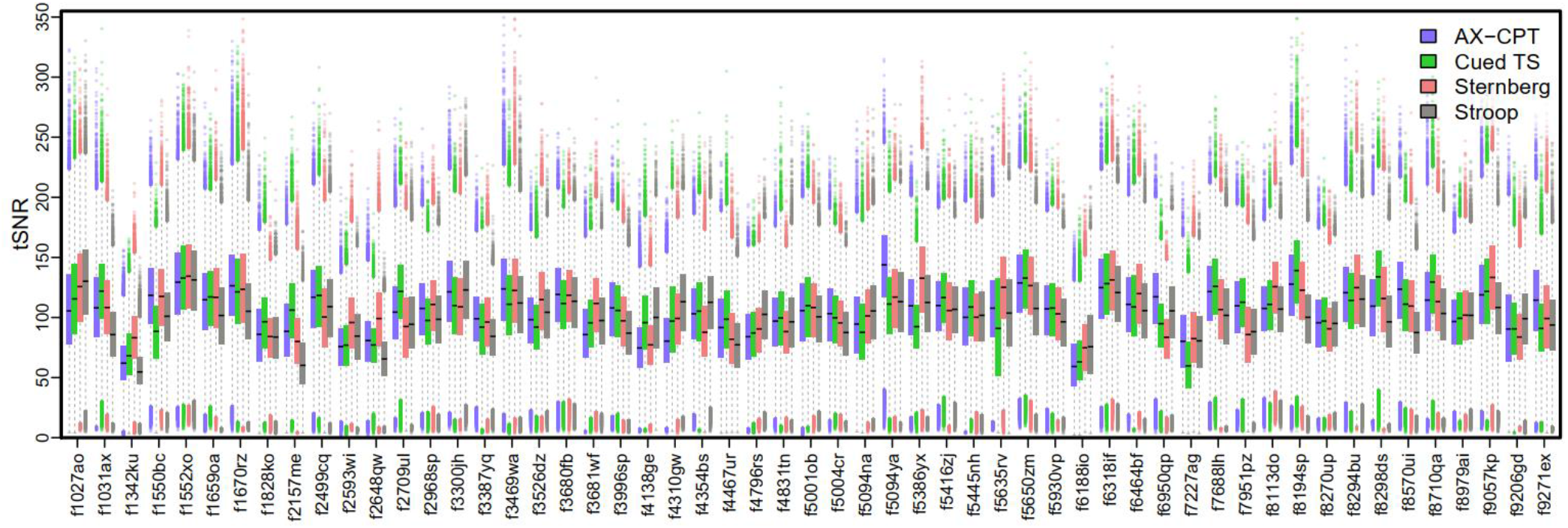
Distribution of tSNR for each subject and task (both runs of each task combined), calculated over all surface vertices (QC_tSNR_summary^18^). Surface and volume tSNR and standard deviation images for each individual are shown the corresponding QC files at the DMCC55B OSF site^18^.

### Positive Control Analyses

An important part of the DMCC quality control procedures is positive control analyses. These check for strong effects that should be present and easily detectable, even within single-subject data, and whose absence suggests poor signal quality or pipeline error. To avoid circularity, control analyses must have high face validity, and be independent of experimental hypotheses. The use of positive control analyses is especially valuable in task fMRI datasets, given the lack of unambiguous methods for estimating the relevant contrast to noise ratio, or thresholds for motion or tSNR^105–107,111,112^. Currently, we use two positive control analyses for quality control, implemented as separate GLMs. GLMs were chosen since many of the DMCC target analyses involve GLMs, and it is generally advisable to use similar techniques for the experimental and control analyses. However, for illustration purposes, here we present control analyses implemented without GLMs, in order to have the code be as self-contained and easy to run as possible; secondarily, we present GLM-free analyses as a demonstration that when effects are strong they can be detected with minimal modelling.

The first control analysis, “buttons,” defines each button press (only) as an event; the contrast of button presses against baseline should show somatomotor activation. Further, in the DMCC, fingers on the right hand are always used for button presses, so associated somatomotor activation should be most prominent in the left hemisphere. Task responses are often suitable targets for positive control analysis, because the occurrence of movement can be objectively verified (unlike e.g., psychological states, whose existence is necessarily inferred), and motor activity is generally strong, focal, and located in a low *g*-factor area^113^.

Figure 9 shows the results of a “buttons” positive control analysis of the DMCC55B Sternberg task data, implemented as ROI-based multivariate pattern analysis (MVPA)^114^. Briefly, the timecourses in each surface vertex of preprocessed Sternberg run data were normalized and detrended (with afni, 3dDetrend -normalize -polort 2). Next, frames from 3 to 8 seconds after each button press (to approximate the hemodynamic delay) were averaged together to create summary images for each button press, and an equal number of non-overlapping randomly selected frames were averaged together to create no-response images (e.g., for a button press that occurred in frame 26, frames 28 to 33 would be averaged together; frames 15 to 20 could be averaged together for a no-press image; the Sternberg task was used for this analysis because the long time between responses was most conducive to such temporal subsetting). Then, these images were averaged again, yielding one example per class (button press or not), person, and run. Leave-one-subject-out cross-validation (55-fold) was then performed (linear SVM, c=1) separately in each cortical parcel of the 1000-parcel, 17-network Schaefer parcellation^115^. Final classification accuracy (Figure 9) is the average of these 55 folds.

**Figure 9.**
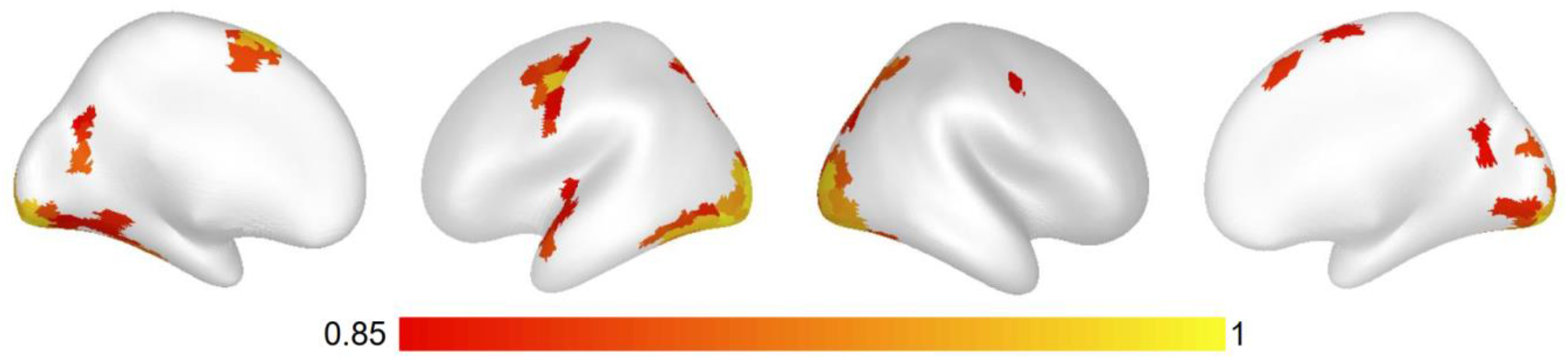
Classification accuracy from a positive control MVPA “buttons” analysis (classifying whether examples were after a button-press or not), Sternberg task. Leave-one-subject-out cross-validation within surface Schaefer 1000×17 parcels^115^, linear SVM (c=1), default scaling, chance is 0.5. Classification accuracy was greatest in visual and left somatomotor parcels, as expected since the right hand was used for button pressing and the stimuli screens varied in appearance between response and baseline periods (Figure 3). Code and additional analyses in the controlAnalysis files^18^.

This control analysis yielded a few parcels with very high (> 90%) classification accuracies (chance 50%; details in the controlAnalyses_buttons files^18^). These parcels were located in the expected anatomic locations (Figure 9), especially left hemisphere somatomotor and bilateral visual areas (most no-button-press events were in ITI or break periods without presented words, producing a pronounced visual difference between the classes; see Figure 3). The anatomic distribution and magnitude of highly-classifying parcels suggests that task-related (here, button presses) information was detected (the positive control analysis was successful), implying that there was not a catastrophic failure in the dataset or processing.

The second control analysis, “on-task” or “ONs”, evaluates general event-related activation by giving all trials the same label; contrasting all trial-related activation against baseline (here, ITIs and the periods before and after the task blocks). For the DMCC tasks these analyses should produce widespread activation in visual, motor, and higher-order cognitive areas (e.g., frontoparietal), both in individuals and at the group level, and so are useful for identifying potential problems with the analysis pipeline or a run’s signal quality. As with “buttons”, failure of the “on-task” analysis suggests an overwhelming error, and that further analysis should not occur until it is resolved.

Figure 10 shows results from an on-task analysis of the Stroop task based on averaging (the short Stroop trials make it the most suitable task for this procedure). Briefly, this analysis also began by normalizing and detrending the timecourse for each surface vertex, then averaging together the timecourses for all vertices within each parcel, yielding one timecourse for each person, parcel, and run. The onset of each trial was used to extract its corresponding timecourse frames, which, averaged over all trials, resembles the canonical HRF in task-responsive parcels (Figure 10a), as expected. The response can further be summarized by averaging the peak frames (here, 3 to 6 seconds after each event), yielding a single number per person, run, and parcel. Figure 10b plots these averages for the group (top) and two representative participants (middle and lower); other participants and details in controlAnalyses_ONs^18^. Overall, these areas resemble those often labelled the task positive and negative networks, as expected, further suggesting that there was not a catastrophic failure in the dataset or processing.

**Figure 10.**
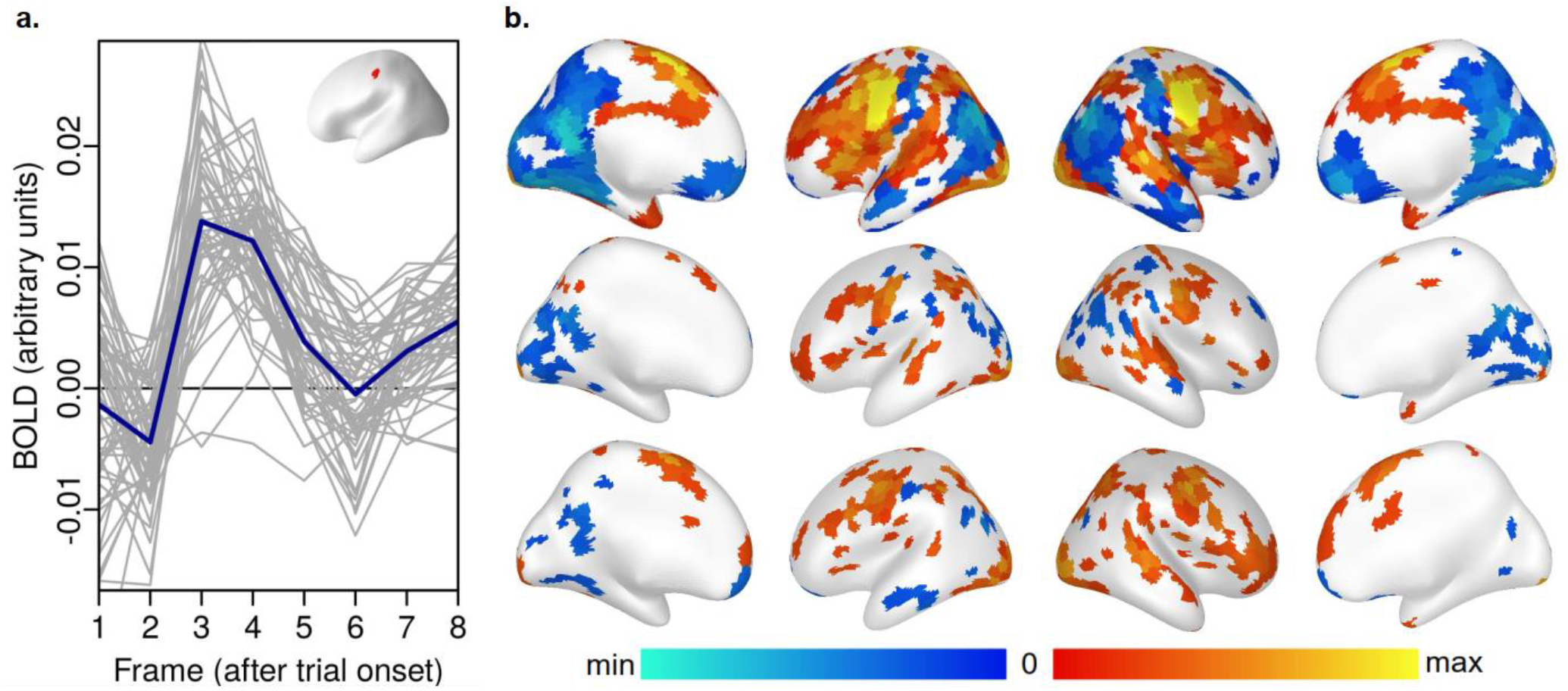
Parcel averages from a positive control “on-task” analysis (average BOLD after trial onsets), Stroop task. Panel a: Example parcel-average timecourses, which resemble the HRF. This is parcel 116, run 1; grey lines are individual participants, blue the group mean. Panel b: Average of frames 2-5 for each parcel; the top row is the group average, while the middle and bottom rows show two individual participants with representative differences from the mean (f5386yx, f5445nh run 1) lower. Code and additional examples in the controlAnalysis files^18^.

## Usage Notes

To maximize reuse, DMCC55B includes a minimum of demographic information, and identifies participants by unique DMCC IDs, rather than those assigned to the same individuals by the HCP or the NIH NDA GUIDs. The DMCC is a Connectome Coordination Facility (CCF) project, and recruits (a subset of) participants who took part in the HCP Young Adult study^11^. The subject key mapping DMCC and HCP IDs is stored in the HCP ConnectomeDB, titled “DMCC (Dual Mechanisms of Cognitive Control) subject key”. Researchers with HCP restricted-level access can obtain this subject key, and must then follow the HCP data usage guidelines for both the mapping and any information collected on the participants in other CCF projects.

## Code Availability

All custom code is freely available in several locations, depending on type. Code to work with the DMCC55B data records, including to reproduce the validation analyses, is in the DMCC55B Dataset Description component^18^ of the DMCC OSF site^15^. Code elsewhere is not specific to the DMCC55B dataset, but used in the main DMCC project. Eprime task presentation scripts can be downloaded after completing the online form^58^. Processing and summary scripts used after data collection are in the dualmechanisms GitHub^69^ or Docker Hub^116^ repositories, depending on type.

## Acknowledgements

We thank Maria Z. Gehred, Erin Gourley, Leah Newcomer, and Kevin Oksanen for assistance with data collection and processing. We thank Nicholas Bloom, Mitch Jeffers, and Carolina Ramirez for assistance with pipeline programming and dataset management. Research funded by NIH R37MH066078 to Todd S. Braver

JAE designed analyses, wrote code and the manuscript. TSB designed the experiment, obtained funding, and wrote the manuscript. All authors contributed to discussions and analyses, and edited the manuscript.

The authors declare no competing interests.

